# Antagonistic, synergistic, and social pleiotropy in microbial cheaters

**DOI:** 10.1101/2022.08.11.503674

**Authors:** Pauline Manhes, Kaitlin A. Schaal, Gregory J. Velicer

## Abstract

Cooperation is widespread among microbes. One mechanism proposed to constrain cheating is antagonistic pleiotropy, wherein mutations that cause defection from cooperation, while potentially under positive selection for this effect, reduce fitness at other traits. The bacterium *Myxococcus xanthus* engages in pleiotropically connected complex multicellular behaviors, including motility, predation, and starvation-induced fruiting-body development. Sporulation during development is susceptible to cheating. Here we investigate pleiotropic impacts on how cheater spores respond to stressful environmental changes after development, relative to cooperator spores. A cheater with a mutation in the developmental-signaling gene *asgB* shows antagonistic pleiotropy under both heat and basic-pH stress, while a cheater mutated at *csgA* shows synergistic pleiotropy under basic-pH stress. Further, in a social form of pleiotropy, cooperator spores formed in mixture with the *asgB* cheater are less resilient under basic pH than those from pure groups; interaction of cooperators with cheaters reduces the cooperators’ physical robustness. Our results indicate that, depending on the mutation, pleiotropy can promote as well as limit cheating alleles. They additionally demonstrate that alleles can pleiotropically alter traits in organisms not carrying those alleles. Synergistic and social pleiotropy may contribute to shaping the evolutionary dynamics of cooperation and cheating in many social systems.

---

Defectors contribute less to cooperative interactions than others (Dugatkin 1990, Turner and Chao 2003). This can generate a fitness advantage from exploitation of others’ cooperative acts, which is often referred to as cheating (Axelrod and Hamilton 1981). Cheating-driven increases of less cooperative genotypes can reduce group productivity (Maclean *et al*. 2010), an effect known as cheating load (Velicer 2003). Concomitantly, selfish behavior can threaten the stability of cooperative systems (Crespi 2001, Dunny *et al*. 2008, Bourke 2011) and, in the extreme, even cause extinction of entire populations (Fiegna and Velicer 2003). Among microbes, cheater mutants have emerged spontaneously in laboratory populations of diverse species (Zambrano et al. 1993; Velicer *et al*. 1998, 2000; Harrison and Buckling 2005; Dunny *et al*. 2008; Jiricny *et al*. 2010; Manhes and Velicer 2011) and are thus expected to arise often in wild populations (Sandoz *et al*. 2007).

Yet many and diverse forms of microbial cooperation thrive (Crespi 2001, West *et al*. 2007), raising the question of which mechanisms contribute most to limiting cheating in any given cooperative system (Fletcher and Doebeli 2009). Many such mechanisms have been proposed (Travisano and Velicer 2004; Griffin *et al*. 2004; Gilbert *et al*. 2007; Strassmann and Queller 2011, Ågren *et al*. 2019), including pleiotropy, in which one genetic locus affects multiple distinct organismal traits (Mayr 1963; Stearns 2010; Wagner and Zhang 2011). Antagonistic pleiotropy in particular, in which an allele enhances fitness at one trait while reducing it at another, has been suggested to play an important role in the evolution of cooperation by limiting the degree of cheating that evolves in cooperative systems (Foster *et al*. 2004, Ostrowski 2019, Jahan *et al*. 2022). Cooperative traits can be promoted when they are linked to other traits, because mutation to a cheating allele has multiple effects (Bentley *et al*. 2022, but see dos Santos *et al*. 2018). Antagonistic pleiotropy has also been invoked to explain the persistence of alleles associated with aging (Williams 1957; Charmantier *et al*. 2006; Magwire *et al*. 2010) and disease (Carter and Nguyen 2011), and suggested to mediate ecological specialization (Cooper and Lenski 2000; MacLean *et al*. 2004).

Antagonistic pleiotropy might limit cheating in two different ways, either by preventing defectors from successfully cheating in the first place or by causing defectors that do cheat on a cooperative trait to have a disadvantage in some other respect. Defection mutations resulting in intrinsic defector inferiority - the failure of defection to generate a cheating advantage (Travisano and Velicer 2004) - may be common, as several cooperation-defective microbial mutants unable to cheat have been identified (Velicer *et al*. 2000, Foster *et al*. 2004, Sathe *et al*. 2019). For example, in *Dictyostelium discoideum*, a eukaryotic microbe that sporulates within a fruiting body (Kessin and Campagne 1992), cells with a mutation in the gene *dimA* migrate disproportionally into the aggregate region where cells normally become spores. However, they are ultimately unable to cheat at sporulation because the *dimA* mutation also diminishes the ability of mutant cells to transform into spores (Foster *et al*. 2004). Defection mutations that do allow cheating but otherwise pleiotropically harm cheaters may also be common (Cameron-Pack *et al*. 2022, Vulić and Kolter 2001, Dandekar *et al*. 2012, Wang *et al*. 2015). For example, in *Pseudomonas aeruginosa*, mutations that allow cheating on siderophore production can decrease the ability of cheaters to form biofilms (Banin *et al*. 2005; Harrison and Buckling 2009).

Despite this evidence, the overall role of pleiotropy in the evolution of cooperation remains largely unresolved, with authors diverging substantially in their view of its importance (Bentley *et al*. 2022, dos Santos *et al*. 2018, Sathe *et al*. 2019, Kramer *et al*. 2019, Figueiredo *et al*. 2021). The direction of influence may indeed be the other way around, with cooperation influencing pleiotropic linkage (dos Santos *et al*. 2018). Our ability to draw general conclusions is hindered by the small number of social organisms, cooperation genes, distinct mutations within a given gene, and pleiotropy-impacted traits which have actually been investigated. Expanding the range of studied social systems can yield surprises, for example the finding that cheaters can exhibit positive frequency dependence of fitness as well as negative (Sathe *et al*. 2019). Moreover, interest in antagonistic pleiotropy as a cheater-limiting mechanism may have overshadowed consideration of how other forms of pleiotropy might impact cooperation.

Here we test whether spores produced by two cheater mutants of the model social bacterium *Myxococcus xanthus* exhibit altered tolerance of environmental stress relative to spores produced by a wild-type cooperator (Fig. 1). We also test for a social form of pleiotropy in which an allele impacts a trait in the organism that carries it and also impacts a trait in a non-carrier via social interaction, which we refer to as ‘social pleiotropy’ for brevity (see Methods for further discussion of this term). Specifically, we test whether interaction between cheaters and cooperators alters the stress tolerance of the cooperators’ spores.

**Figure 1.**
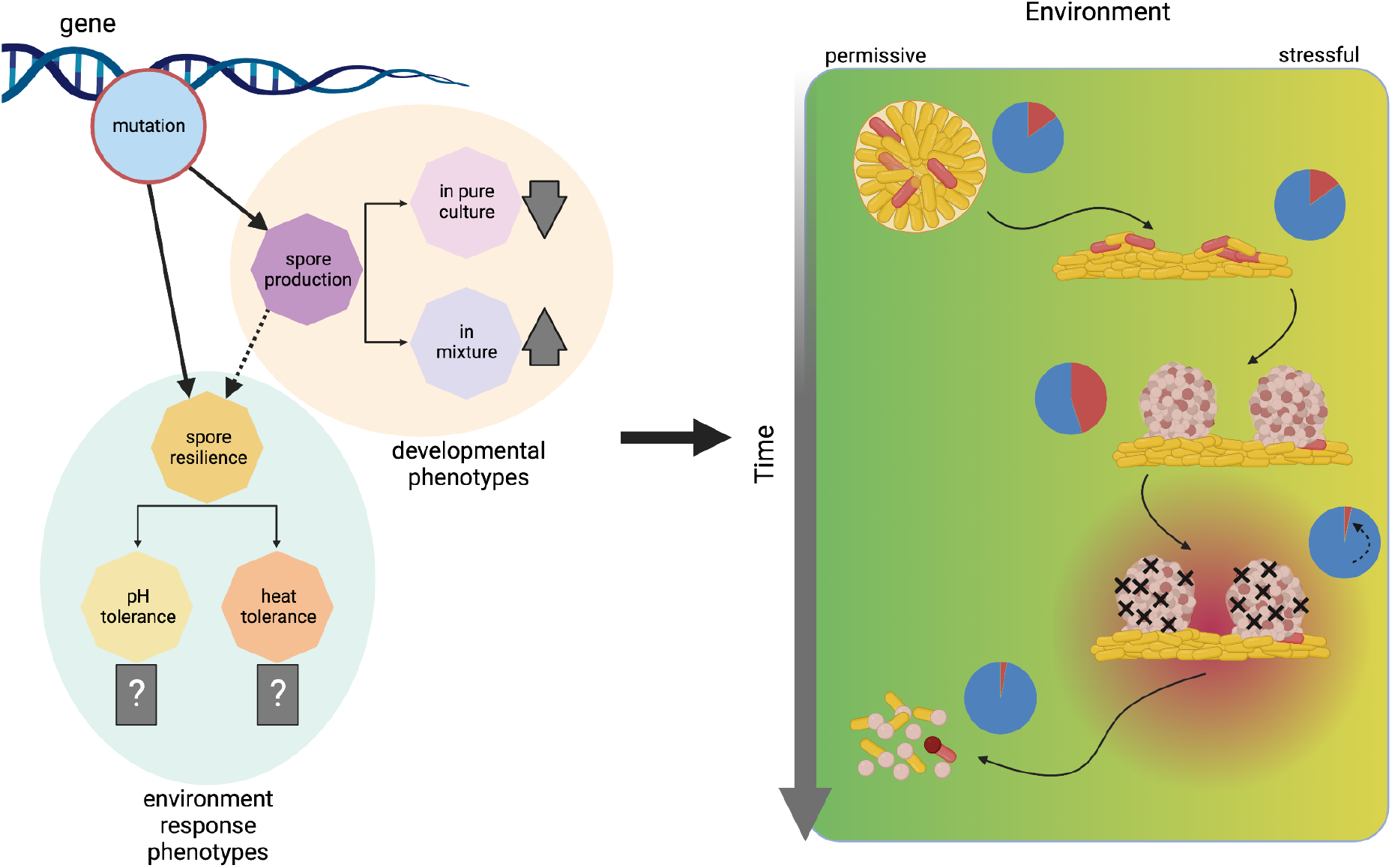
Conceptual overview. Mutations which affect social phenotypes such as starvation-induced sporulation may also impact environment-response phenotypes such as stress tolerance of the spores (left side). These pleiotropic effects may manifest as the environment changes over time, dynamically influencing the ratio of cheaters (red bacteria) and cooperators (golden bacteria) in a mixed population (right side). Here, environmental quality is indicated by color (green = high, yellow = poor, red = exceptionally stressful) and the ratio of cheaters to cooperators is shown in the pie charts (red = cheaters, blue = cooperators). In the case pictured, *M. xanthus* undergoes multicellular development as environmental quality declines due to lack of nutrients, and developmental cheaters increase in frequency during this phase. Later, onset of an exceptionally stressful environment interacts with a pleiotropic phenotype of the cheater spores, causing them to perish at a higher rate than the cooperator spores. By the time the environment improves, the fitness gain from cheating has been lost and the cheaters have declined overall. This scenario constitutes antagonistic pleiotropy; however, synergistic pleiotropy is also possible, in which the pleiotropic phenotype of the cheater spores causes them to perish at a *lower* rate than the cooperator spores and thus augment their advantage gained from cheating. Created with BioRender.com.

Like *D. discoideum, M. xanthus* aggregates upon starvation to form multicellular fruiting bodies composed of tens of thousands of cells, a subset of which transforms into stress-resistant spores (Shimkets *et al*. 2006). Several extracellular signals mediate these processes, including the A- and C-signals (Shimkets 1999). The A-signal is a mixture of amino acids that functions in the beginning of development as a quorum-sensing signal and requires the action of the putative transcription factor *asgB* (Kuspa *et al*. 1992, Bretl and Kirby 2016). The exact nature of C-signaling remains controversial (Kroos 2017), but previous studies have demonstrated a key role of *csgA* (Shimkets and Rafiee 1990, Konovalova *et al*. 2012, Boynton and Shimkets 2015).

Strains lacking either signal cannot form normal fruiting bodies or sporulate well by themselves (Kuspa *et al*. 1986; Kruse *et al*. 2001; Boynton and Shimkets 2015). Both *asgB* and *csgA* mutants can cheat in chimeric groups by exploiting signals produced by the wild type and converting a higher proportion of their cells into spores (Velicer *et al*. 2000). Here we ask whether spores produced by *asgB* and *csgA* mutants exhibit different levels of resilience against environmental stresses than do wild-type spores. We test for differences in stress tolerance of cheater and cooperator spores under two naturally occurring types of stress: pH and heat. *M. xanthus* resides in a wide variety of soil habitats worldwide (Dawid 2000) that can vary greatly in pH (Dimitriu and Grayston 2010), an environmental factor to which vegetative *M. xanthus* cells are highly sensitive (Fiegna *et al*. 2021). Additionally, *M. xanthus* spores produced by different developmentally proficient natural isolates show variation in heat resistance (Kraemer *et al*. 2010). These traits may thus be relevant for understanding the physical robustness of spores with respect to natural stressors.

## Material and Methods

### Bacterial strains

We used as our cooperative wild-type strain GJV2 (Fiegna *et al*. 2006), a variant of the lab type strain GJV1 (Velicer *et al*. 2006) (aka strains R and S, respectively, in Velicer *et al*. (1998)) with a spontaneous mutation in *rpoB* that confers resistance to rifampicin (Zee *et al*. 2014). Our mutant strains were DK4324 (Kuspa *et al*. 1986, Mayo and Kaiser 1989) and DK5208 (Shimkets and Asher 1988, Kashefi and Hartzell 1995, Velicer *et al*. 2000, Kruse *et al*. 2001, Manhes and Velicer 2011, Schaal *et al*. 2022), which have loss-of-function mutations in the A-signal gene *asgB* and the C-signal gene *csgA*, respectively. Mutants defective at these loci were previously shown to produce very few spores in monoculture relative to wild-type, while in mixes with the wild-type the mutants produce more spores than the wild-type, *i*.*e*. they “cheat” (Velicer *et al*. 2000). DK4324 is resistant to kanamycin whereas GJV2 is not. We grew strains in CTT liquid (Bretscher and Kaiser 1978) at 32 °C and 250 rpm.

### Development assay

We initiated development as in Fiegna and Velicer (2003). We grew strains to mid-exponential phase before resuspending in TPM (Kroos *et al*. 1986) to a density of ∼5 × 10^9^ cells/ml. In mixed culture competitions, we combined mutant strains with GJV2 at a 1:9 ratio to the same total density. We inoculated 100 µl of bacteria into the center of TPM 1.5% agar plates and allowed development to proceed for 3 days at 32 °C before harvesting.

### Post-development stress treatments

Harvesting is typically followed by 2 hrs of heating at 50 °C to kill vegetative cells and thus ensure that CFU counts quantify the spore population alone (Velicer *et al*. 2000). For heat treatments, we harvested spores into 1 ml ddH_2_O and incubated for 2 hrs at 50 °C, 55 °C, or 58 °C. For pH treatments, we harvested the spores into 1 ml buffer: phosphate-citrate buffer pH 5.1, Tris HCl pH 6.8 (control for standard conditions), or Tris HCl pH 8.7 (all pHs +/- 0.2) and incubated at room temperature for 18 hours, then at 50 °C for 2 hrs. For all treatments, we then sonicated the spores, diluted them, and plated them into selective and nonselective CTT 0.5% agar. For DK4324:GJV2 mixes, we added 40 µg/ml kanamycin to count CFUs of DK4324. For DK5208:GJV2 mixes, we added 5 µg/ml rifampicin to count CFUs of GJV2. We estimated the spore production of paired strains by subtraction of counts on selective agar from total counts on non-selective agar.

We photographed fruiting bodies (Fig. S1) of GJV2, DK4324, and the mix of the two with an Olympus DP80 camera mounted on an Olympus SZX16 microscope, captured using cellSens software version 1.15 (Olympus, Tokyo, Japan).

### Stress tolerance and relative fitness

We calculated the stress tolerance of the spores produced by each strain by comparing their post-development survival under higher temperatures or acidic/basic pH to their post-development survival under standard conditions. Stress tolerance for strain X is given as

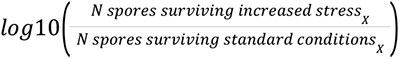

For example, a stress tolerance value of -2.0 indicates that the number of viable spores surviving in a non-standard environment was 1% of the number surviving standard conditions.

We tested for relative fitness differences between the cheaters and GJV2 in mixed competitions with the previously-published parameter *W*_*ij*_ (Vos & Velicer 2009, Schaal et al. 2022):

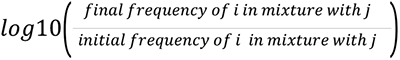

where *i* is the mutant strain and *j* is the cooperator. If *W*_*ij*_ is positive, then *i* is said to cheat on *j*, showing a fitness advantage based on an increase in frequency in mixed competitions, in contrast to its pure-culture defect.

In 13 instances, we estimated a maximum undetectable spore count because no colonies grew even from the lowest dilution. This approach was only necessary when survival was much lower under stressful conditions than under standard conditions and represents a conservative estimate of the effect of environmental stress. In the four cases where we counted zero spores at the lowest dilution for both the cheater and the cooperator in mixture, we used a maximum undetectable count when analyzing the effect of treatments on raw spore counts but excluded these cases from calculation of our metrics of stress tolerance and relative fitness.

### Statistical analyses

We performed all analyses and data visualization in R version 4.2.1 (R Core Team 2018) and RStudio version 2022.07.0 (RStudio Team 2020). We calculated general linear models with environmental level and bacterial strain as fixed factors, and we performed post-hoc Tukey HSD tests, *t*-tests, or Dunnett tests to check specific contrasts. All cases of multiple *t*-tests were corrected using the Bonferroni-Holm method. All data and statistical analyses are available on Dryad (doi: TBD).

### Semantics

In this study, we use ‘social pleiotropy’ to refer to a scenario where an allele impacts one trait in a non-carrier (*e*.*g*. spore robustness) through interaction with the carrier, in which the allele also impacts a different trait (*e*.*g*. spore productivity). A related term used previously is ‘group-level pleiotropy,’ used for example to refer to alleles that impact fitness of non-carriers through group-level effects (Woodley of Menie *et al*. 2017). ‘Social pleiotropy’ has also been used in the context of social animals to indicate when an individual trait impacts “several features of the social organization” (Thierry 2004) or when one locus impacts multiple social traits (Fouks *et al*. 2016). We emphasize that pleiotropy can be considered social in at least two different ways: when an allele impacts multiple distinct social traits of the organism carrying that allele, or when an allele impacts different traits in carriers and non-carriers, with the latter brought about by interaction with the carrier.

## Results

### Cheating under standard conditions

As expected, both mutants showed developmental defects in pure groups that resulted in very few viable spores: ∼0.03% and ∼0.0002% of GJV2 spore number for DK4324 and DK5208, respectively (Fig. S2; Tukey tests for difference from GJV2, *p* < 0.002 for both mutants). DK4324 produced more spores than DK5208 (same Tukey tests as above, *p* = 0.024).

As expected from previous findings (Velicer *et al*. 2000, Schaal *et al*. 2022), both mutants exhibited cheating phenotypes in mixed cultures with GJV2 under standard developmental conditions, converting a larger proportion of their cells into viable spores than did GJV2. DK4324 clearly cheated on GJV2 under the standard conditions used for the pH-treatment comparisons (Fig. 2, pH 6.8; paired one-tailed *t*-test for difference of *W*_*ij*_ from zero, *p* = 0.026) and showed a strong suggestion of cheating in the standard treatment for the temperature comparisons (Fig. 2, 50 °C; same *t*-tests as above, *p* = 0.061). (The pH-control treatment differed from the temperature control treatment only in (i) the additional step of keeping the spores in liquid suspension for 18 hours before heating and (ii) the use of pH 6.8 Tris HCl buffer solution rather than double-distilled water during heating and sonication.) The sporulation superiority of DK5208 over GJV2 in mixture was very clear in both the pH and temperature standard treatments (Fig. 2, purple labels; same *t*-tests as above, *p* < 0.0005 in both cases).

**Figure 2.**
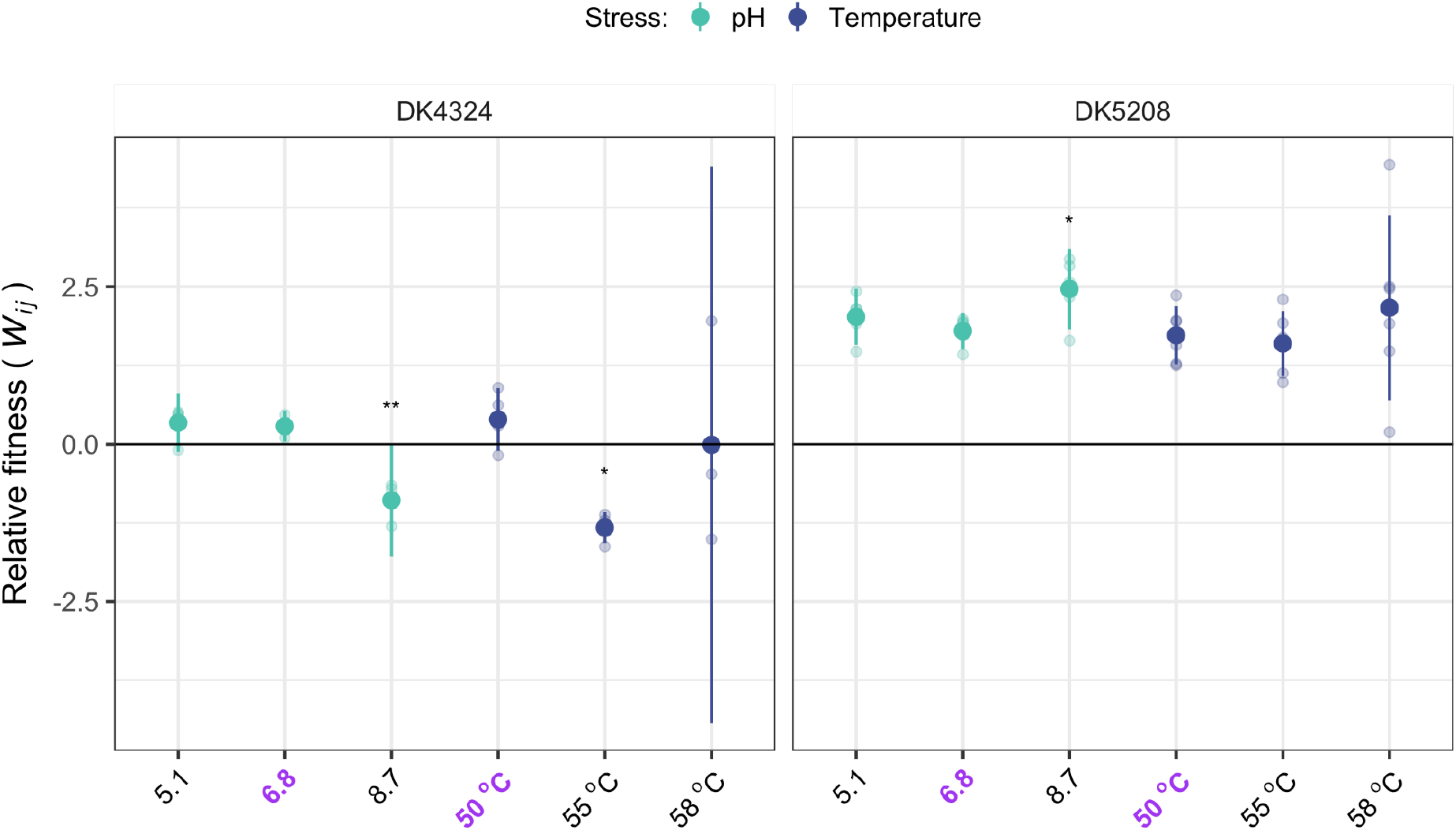
Post-development stress can both eliminate and augment cheater fitness advantages. We calculated the relative fitness of each cheater in comparison to the cooperator in mixture, parameter *W*_*ij*_, as described previously. Positive and negative values indicate that the cheater produced proportionally more or fewer spores that survived the specified post-development treatment than the cooperator. Small transparent circles show individual replicates, and large solid circles show averages. Bold purple labels indicate standard treatments. Error bars are 95% confidence intervals. 3-6 biological replicates. Asterisks indicate a significant difference from the respective standard conditions (purple label): * *p* < 0.05, ** *p* < 0.01.

### Stress effects of treatment conditions

Additionally, we subjected spores produced by the cooperator or by mixtures of the cooperator and a defector during development to different pH and temperature environments. Strains generally found the non-standard conditions to be more stressful than the standard conditions, *i*.*e*. spore survival was less than under standard conditions of pH 6.8 for the pH treatments and of 50 °C for the temperature treatments. (Fig. S3; two-tailed paired *t*-tests for difference in spore count from standard conditions for cooperator in pure culture, cooperator in mixture, and defector in mixture, *p*-values < 0.026). The slight exception is pH 5.1, where a lower survival rate of GJV2 spores in pure culture compared to pH 6.8 is not quite significant (same *t*-tests as above, *p* = 0.054), and where neither of the defectors (same *t*-tests as above, *p*_DK4324_ = 0.14, *p*_DK5208_ = 0.36) nor GJV2 in mixture with DK4324 (same *t*-tests as above, *p* = 0.16) experienced a reduction in spore survival. This result might be viewed as surprising given that acid stress at pH 5.1 has been found to kill intermediate- and low-density cultures of vegetative *M. xanthus* cells (Fiegna *et al*. 2021).

Interestingly, DK4324 spores appear to be more sensitive to stress than DK5208 spores. DK4324 spore survival in mixture decreased more both under 55 °C conditions and under basic pH stress than did DK5208 spore survival in mixture (Figs. 3, 4; *t*-tests for differences in stress tolerance between cheaters across all stress treatments, *p* = 0.007 for both). For 58 °C and pH 5.1 conditions, we observed no difference between the two cheaters (same *t*-tests as above, *p* = 0.3 for both).

**Figure 3.**
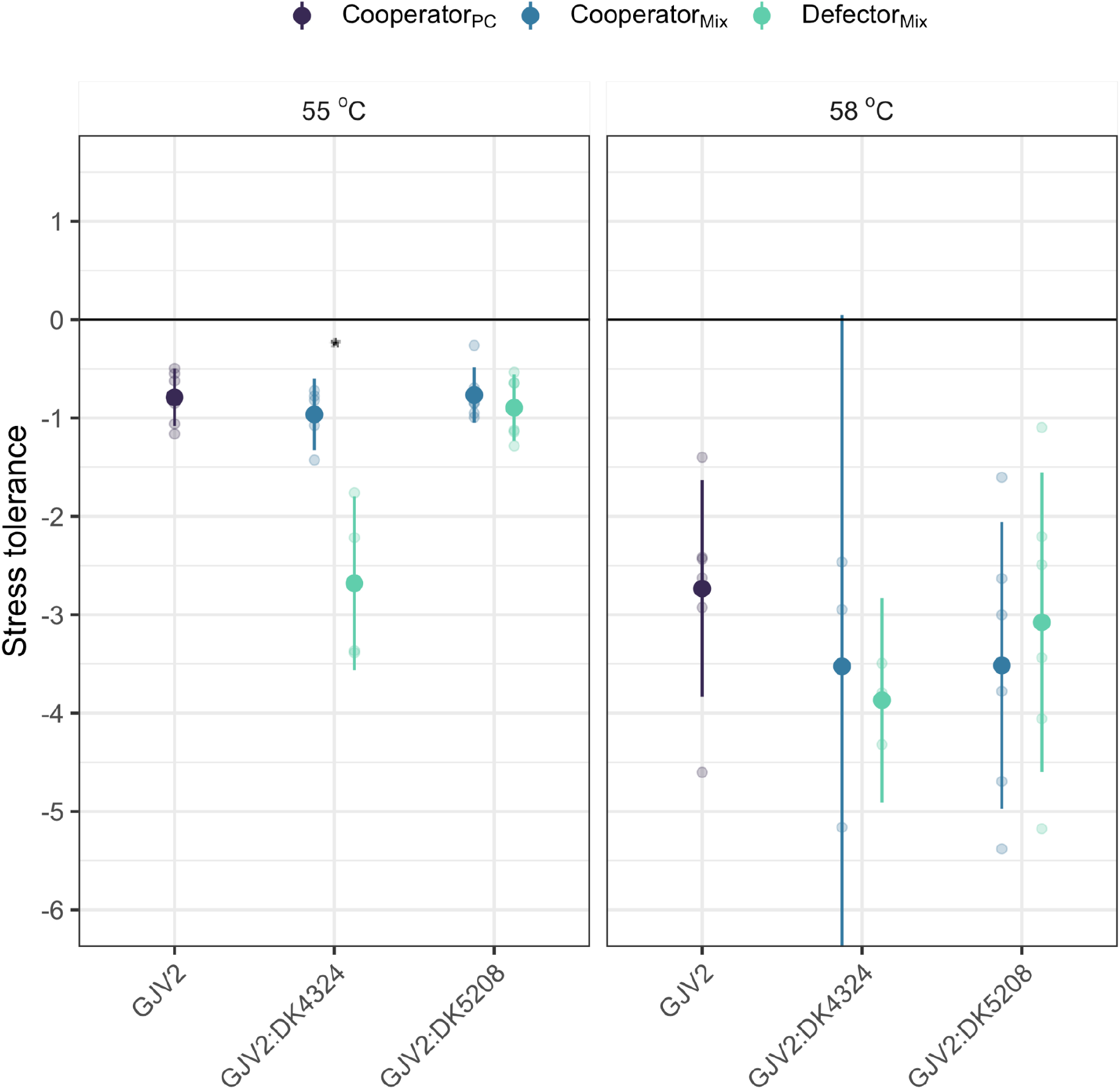
DK4324 spores are uniquely sensitive to increased post-development heat stress. We calculated the stress tolerance of spores of each strain as the log_10_ of the number of viable spores after surviving increased heat stress (55 or 58 °C) relative to the number of viable spores surviving standard spore-selection conditions (50 °C). For example, a value of -1 indicates that spore survival under increased stress is 10% of that under standard conditions. ‘Cooperator’, strain GJV2; ‘Defector’, strains DK4324 and DK5208; ‘PC’, pure culture; ‘Mix’, mixed culture with a 9:1 cooperator:defector ratio at the onset of starvation. Small transparent circles show individual replicates, and large solid circles show averages. Error bars are 95% confidence intervals. The asterisk indicates a significant difference between the stress tolerance of GJV2 and DK4324 in mixed culture. 3-6 biological replicates.

**Figure 4.**
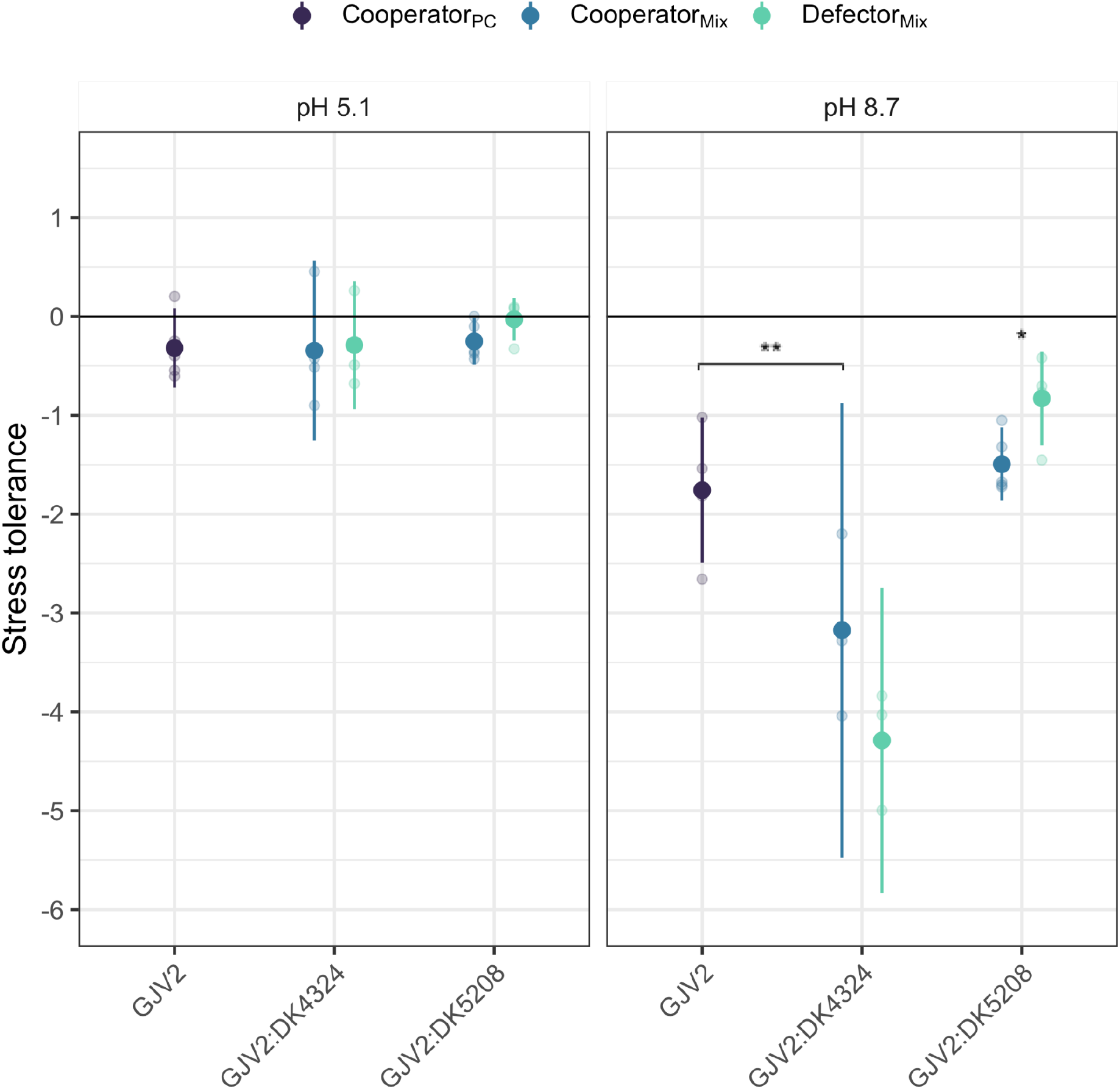
DK5208 spores are uniquely resilient against post-development pH stress; interaction with the *asgB* mutant decreases cooperator spore resilience. The stress tolerance of spores of each strain is calculated as the number of viable spores (log_10_) that survived post-development pH stress relative to the number surviving under standard conditions (pH 6.8). Thus, a value of -1 indicates that spore survival under the specified stress was 10% of that under standard conditions. ‘Cooperator’, strain GJV2; ‘Defector’, strains DK4324 and DK5208; ‘PC’, pure culture; ‘Mix’, mixed culture with a 9:1 cooperator:defector ratio at the onset of starvation. Small transparent circles show individual replicates, and large solid circles show averages. Error bars are 95% confidence intervals. Asterisks indicate significant differences between the stress tolerance of GJV2 in pure culture vs. GJV2 in mixture with DK4324 (left) and between GJV2 vs. DK5208 in mixed culture (right) (* *p <* 0.05, ** *p* < 0.01). 3-5 biological replicates.

### Antagonistic pleiotropy under increased heat stress

We determined whether the defector mutations had pleiotropic effects on spore hardiness by comparing the stress tolerance of the cheater in mixture to the stress tolerance of the cooperator in the same mixture or by comparing *W*_*ij*_ of the cheater under stress to its *W*_*ij*_ under standard conditions. If a cheater mutation causes cheater spores to differ from wild-type spores in their survival under increased post-development stress (whether heat or pH) relative to their survival under standard conditions, this is evidence of pleiotropy. In mixed cultures, we found that increasing heat from 50 to 55 °C resulted in DK4324 spores surviving the stress at a rate only 2% that of GJV2 (Fig. 3; paired *t*-tests for difference from cooperator, *p* = 0.007). These results indicate that wild-type spores formed in mixed groups are more resilient to heat stress than are DK4324 spores. This lower resilience of DK4324 decreased its post-heat fitness compared to standard conditions (Fig. 2; Tukey tests, *p* = 0.023), and the mutant has an outright fitness disadvantage relative to GJV2 (two-tailed *t*-tests for all *W*_*ij*_’s difference from zero, *p* = 0.0004). At 58 °C, survival of both DK4324 and GJV2 was reduced relative to standard conditions by a factor of ∼10^4^. DK5208 and wild-type spores did not differ in their tolerance of either level of heat stress (same *t*-tests for difference from cooperator as above, *p*’s > 0.56). Moreover, variation in stress tolerance was also higher at 58 °C than at 55 °C (Fig 3; two-tailed *t*-tests for difference in variance of each sample at one stressful condition vs the other, *p* = 0.046). We did not observe such a difference for the pH treatments (Fig 4; same *t*-tests as above, *p* = 0.16).

### Synergistic and antagonistic pleiotropy under basic pH stress

We also compared the effect of pH stress on post-development survival of cheater vs wild-type spores. Incubation at pH 5.1 prior to standard processing appears to have only slightly reduced spore survival across all three genotypes, in both pure-culture and mixed-culture assays (Figs. 4, S3). Incubation at pH 8.7, however, strongly reduced spore survival of all three genotypes in mixed culture and of the wild-type in pure culture (Figs. 4, S3). The low stress tolerance of DK4324 spores at pH 8.7 translated into decreased post-stress relative fitness compared to standard conditions (Fig. 2; Tukey tests, *p* = 0.0012), which eliminated the cheating advantage of this strain seen at pH 6.8.

The *csgA* mutation in DK5208 exhibited a positive pleiotropic effect on this strain’s tolerance of basic-pH stress (paired *t*-tests for difference from cooperator, *p* = 0.028): survival was reduced by 84%, compared to 97% survival reduction of the wild-type. This positive pleiotropy on post-development spore resilience enhanced DK5208’s fitness after basic-pH stress above its cheating advantage exhibited under standard conditions (Fig. 4; Tukey tests, *p* = 0.045).

### Interaction with a cheater reduced the hardiness of wild-type spores

Finally, we compared the stress tolerance of the cooperator in mixture to the stress tolerance of the cooperator in pure culture to determine whether mixture with the defector had a social effect on cooperator-spore hardiness. Most of our treatments of increased post-development stress had similar effects (or non-effects) on the survival of wild-type spores regardless of whether wild-type underwent development in pure culture or in mixture with a cheater (Figs. 3 and 4) but there was one exception to this trend. The wild-type showed lower stress tolerance at pH 8.7 when mixed with DK4324 than in pure culture (Fig. 4; Dunnett test for difference from GJV2 in pure culture *p* = 0.015). Under basic stress, the *asgB* mutation appears to reduce not only the hardiness of spores bearing this mutation but also that of spores formed by a social partner, as well as affecting the morphology of the chimeric fruiting bodies (Fig. S1).

## Discussion

Mutations creating obligate social cheaters provide a fitness advantage in social environments that allow effective exploitation of cooperator genotypes, but decrease fitness in social environments in which exploitation is either not possible (*i*.*e*. pure groups of cheaters or groups with non-compatible cooperators (Schaal *et al*. 2022)) or is insufficient to generate a relative fitness advantage (*e*.*g*. in groups with low cooperator frequencies (Velicer *et al*. 2000, Velicer and Vos 2009)). However, beyond this, social-gene mutations may frequently have pleiotropic effects on mechanistically distinct traits, and whether selection will promote maintenance or loss of a mutation in the long run depends on its net fitness effect across all traits it impacts over time (Otto 2004).

Here we tested whether mutants that are defective at pure-culture spore production and can cheat on cooperators experience pleiotropic effects on stress tolerance. We allowed mixed cultures of cooperators and mutants to undergo development, during which the mutants cheated, then applied environmental stresses which may counteract or enhance the effect of cheating by differentially impacting the two genotypes and thereby influencing relative frequencies. We find that distinct cheating mutations in the same organism can be associated with both negative and positive effects at another trait seemingly unrelated to cheating, namely spore hardiness (Table 1). Examination of only two cheater mutants reveals not only reduced fitness of cheater spores under increased stress (DK4324 under heat and basic-pH stress) but also increased fitness (DK5208 under basic-pH stress). For cheating mutations which confer both synergistic and antagonistic pleiotropic effects, the former may counteract the latter and thus reduce their role in limiting cheater equilibrium frequencies (Van Dyken *et al*. 2011). In another form of synergistic pleiotropy, a single mutation might create “supercheats” with the ability to cheat on multiple social traits (Harrison and Buckling 2009).

**Table 1.** Summary of stress effects on spore survival.

| Cheater genotype | pH stress |  | heat stress |  |
| --- | --- | --- | --- | --- |
|  | pH 5.1 | pH 8.7 | 55 °C | 58 °C |
| DK4324 |  | - | - |  |
| DK5208 |  | + |  |  |

Symbols indicate that antagonistic (-) or synergistic pleiotropy (+) was detected for stress tolerance and/or relative fitness.

Pleiotropy is often conceived of as affecting different traits in the same organism, but our findings illustrate that pleiotropy can also affect a trait in a social partner. Specifically, interaction of the wild-type with the mutant DK4324 reduced post-development survival of wild-type spores under basic pH stress relative to wild-type spores in monoculture. Despite being in the minority, DK4324 visibly altered the fruiting bodies produced in mixture with GJV2 compared to those GJV2 produced in monoculture (Fig. S1). It has been speculated that spores which differentiate within well-formed fruiting bodies may often be more robust than spores in defective fruiting bodies or outside a fruiting body (La Fortezza and Velicer 2021). The negative impact of basic-pH stress on this mixture could thus be due to smaller size or lower structural integrity of the fruiting bodies. This result shows that cheaters can impact cooperator fitness at traits not obviously related to a focal cooperative trait and in ways only manifested beyond the context of a focal cheating interaction, e.g. post-development spore resilience in the case of *Myxococcus*. Future theoretical and experimental investigations of the evolution of cooperation should account for the possibility of such social pleiotropy across distinct life-history stages and environments.

More than two decades of research have yielded many insights into ecological and evolutionary dynamics of cheating and cooperation in microbes (West *et al*. 2021), yet we are far from a thorough understanding of the roles of antagonistic, synergistic, and social pleiotropy in these dynamics. This will require analysis of cheating-allele effects in a wide range of social systems, across a large sample of cheatable social genes, and across many fitness-relevant traits. Species might differ significantly in their relative frequencies of antagonistic versus synergistic pleiotropy of cheating mutations. Mutations in different cooperation genes in the same organism might generate pleiotropy in opposite directions, as exemplified by our two mutants having opposite effects on relative fitness under basic stress. Finally, the same cheating mutation might have both negative and positive effects across different traits in the same organism.

Outcomes of social interactions between defectors and cooperators may depend not only on the specifics of genetic architecture but also on broader impacts of environmental conditions that may shift over ecological and evolutionary time (Fig. 1). Natural populations of microbes experience daily and seasonal shifts in relevant parameters such as temperature, pH, and humidity, and they are increasingly understood to be affected by climate change (Thomas *et al*. 2017; O’Donnell *et al*. 2018; Franco *et al*. 2019; Chase *et al*. 2021; Beaufort *et al*. 2022). Here, we find that increasing the temperature to which spores are exposed can not only cause a cheater to have lower fitness than a cooperator under the increased stress (Fig. 2, DK4324 fitness at 55 °C compared to 50 °C), but can also increase the variation in spore-survival outcomes across replicates for both cooperators and cheaters (Fig. 3, increase from 55 °C to 58 °C for both mutants). The higher temperature was more stressful to spores, resulting in lower survival as well as increased variance. Exposure to pH 8.7 also generally resulted in lower spore survival than to pH 5.1, although the concomitant increase in variation is not significant. However, these results suggest that as environments change, greater stress may result in less-predictable outcomes of interactions, magnifying the role of chance in social evolution. Environmental change may therefore reduce the effectiveness of some evolved cheater-control mechanisms and destabilize some cooperative systems.

## Figures

**Figure S1.**
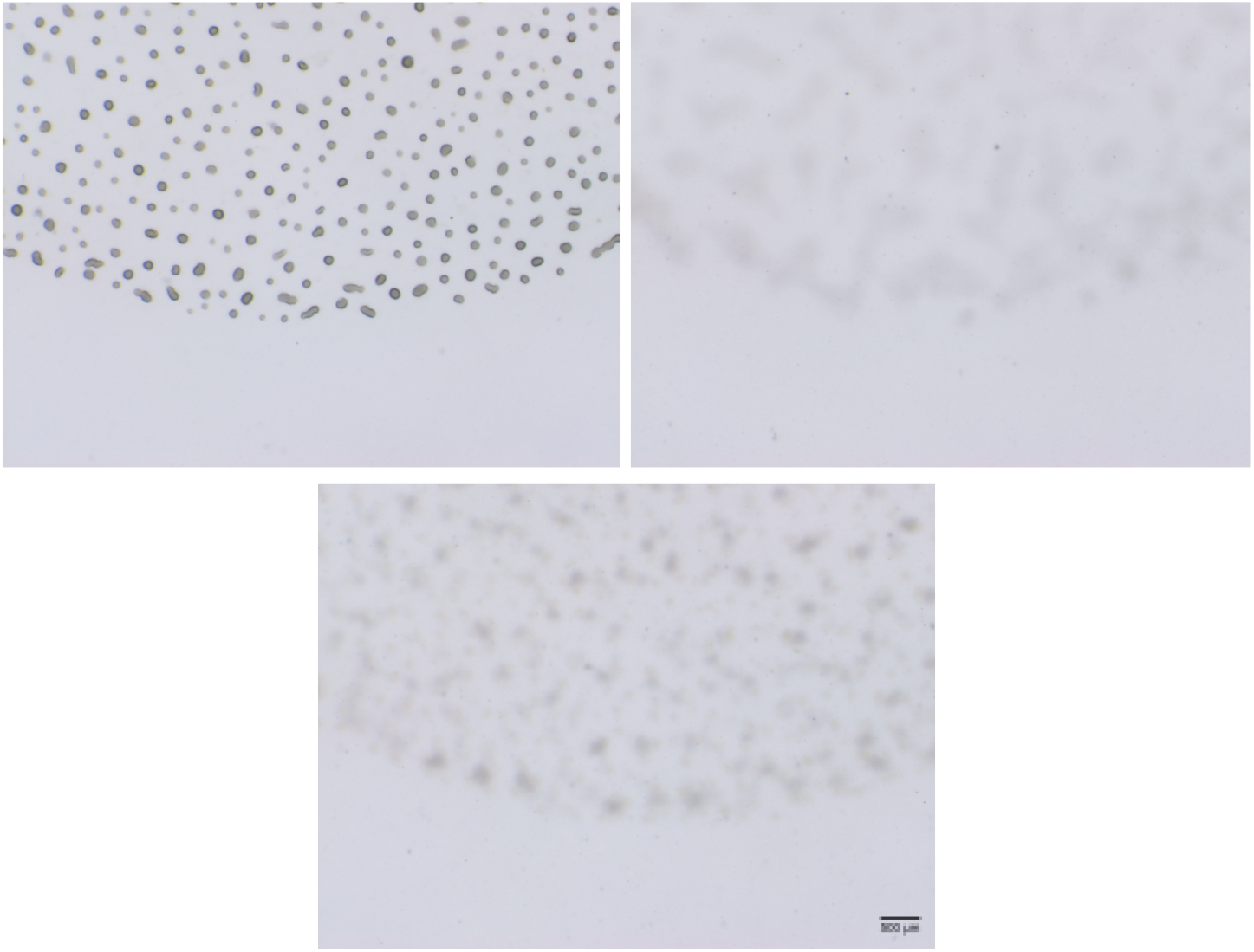
Presence of the *asgB* mutant at only 10% in a mixed group with the cooperator reduces fruiting body quality. Clockwise from the upper left: GJV2 in pure culture, DK4324 in pure culture, GJV2:DK4324 9:1 mixed culture.

**Figure S2.**
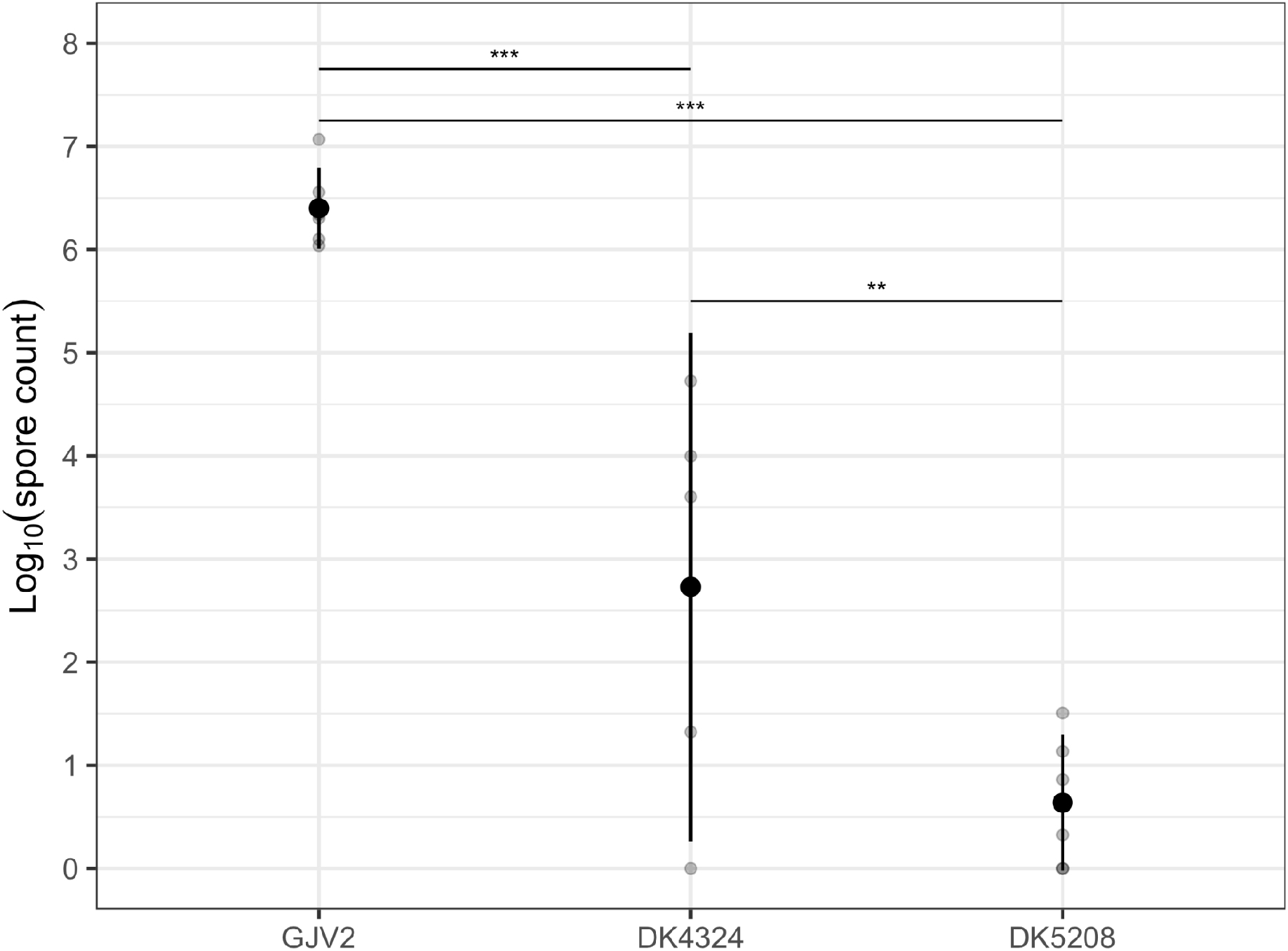
Spore production of each strain in clonal culture under standard lab conditions. Pure-culture spore production of all strains is shown. Spores were selected after development at 50 °C and pH 7.0 in the absence of antibiotics. GJV2 sporulates proficiently, while the mutants show greatly reduced spore production. Small grey circles show individual replicates, and large black circles show averages. Error bars are 95% confidence intervals. 5-6 biological replicates. Asterisks: ** *p* < 0.01, *** *p* < 0.001.

**Figure S3.**
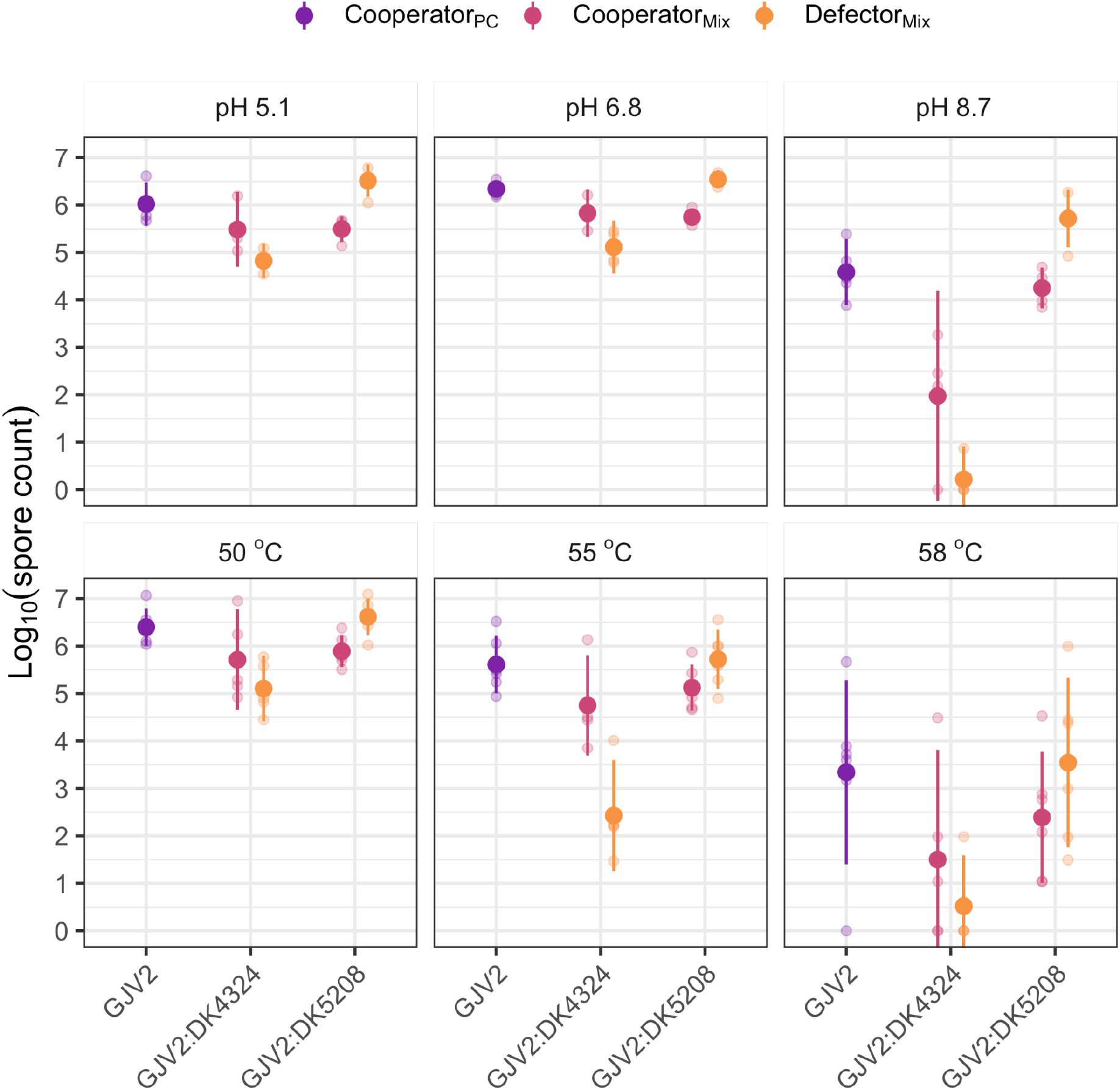
Environmental changes can be stressful to spores. We measured spore production by the cooperator in pure culture (PC) and by both the cooperator and defectors in pairwise mixes under standard conditions (pH 6.8, 50 °C) and under different pHs or increased temperatures to test whether these changes impacted spore survival. Small transparent circles show individual replicates, and large solid circles show averages. Error bars are 95% confidence intervals. 5-6 biological replicates.

